# Robust induction of functional astrocytes using NGN2 expression in human pluripotent stem cells

**DOI:** 10.1101/2022.09.07.507028

**Authors:** Martin H. Berryer, Matthew Tegtmeyer, Loïc Binan, Vera Valakh, Anna Nathanson, Darina Trendafilova, Ethan Crouse, Jenny Klein, Daniel Meyer, Olli Pietiläinen, Francesca Rapino, Samouil L. Farhi, Lee L. Rubin, Steven A. McCarroll, Ralda Nehme, Lindy E. Barrett

**Author notes:** Equal contribution (Martin H. Berryer, Matthew Tegtmeyer).

## Abstract

Astrocytes play essential roles in normal brain function, with dysfunction implicated in diverse developmental and degenerative disease processes. Emerging evidence of profound species divergent features of astrocytes coupled with the relative inaccessibility of human brain tissue underscore the utility of human pluripotent stem cell (hPSC) technologies for the generation and study of human astrocytes. However, existing approaches for hPSC-astrocyte generation are typically lengthy, incompletely characterized, or require intermediate purification steps, limiting their utility for multi-cell line, adequately powered functional studies. Here, we establish a rapid and highly scalable method for generating functional human induced astrocytes (hiAs) based upon transient Neurogenin 2 (NGN2) induction of neural progenitor-like cells followed by maturation in astrocyte media, which demonstrate remarkable homogeneity within the population and across 11 independent cell lines in the absence of additional purification steps. These hiAs express canonical astrocyte markers, respond to pro-inflammatory stimuli, exhibit ATP-induced calcium transients and support neuronal maturation *in vitro*. Moreover, single-cell transcriptomic analyses reveal the generation of highly reproducible cell populations across individual donors, most closely resembling human fetal astrocytes, and highly similar to hPSC-derived astrocytes generated using more complex approaches. Finally, the hiAs capture key molecular hallmarks in a trisomy 21 disease model. Thus, hiAs provide a valuable and practical resource well-suited for study of basic human astrocyte function and dysfunction in disease.

## INTRODUCTION

Astrocytes are the most abundant cell type in the human brain. They play crucial roles in regulating neuronal development, maturation, and synaptic connectivity (Allen and Eroglu, 2017; Sofroniew and Vinters, 2010). Astrocyte dysfunction and defective astrocyte-neuron interactions have been implicated in a wide variety of disorders, including psychiatric, neurodevelopmental and neurodegenerative disorders (Allen and Eroglu, 2017; Pietilainen, 2021; Sofroniew and Vinters, 2010). Astrocytes also play an important role in regulating the cerebral microenvironment by interacting with endothelial and microglial cells that participate in the blood brain barrier (Abbott et al., 2006; Cucullo et al., 2002; Goldstein, 1988; Liu et al., 2020) and communicating with oligodendrocytes via direct contact and secretion of cytokines and chemokines (Amaral et al., 2013; John, 2012; Magnotti et al., 2011).

While brain cell types are largely thought to be conserved across species, an increasing number of studies have uncovered divergent molecular, structural, and functional features of glia. For example, transcriptional comparisons between human and rodent have revealed greater differences in glial gene expression signatures compared with neuronal-associated transcripts, suggesting that glial genes may be evolutionarily less conserved than neuronal genes (Hawrylycz et al., 2012). Moreover, while mammalian astrocytes respond to glutamate and ATP by increasing intracellular calcium concentrations, human astrocytes support different calcium wave dynamics as compared with rodent astrocytes (Han et al., 2013; Oberheim et al., 2006) which has the potential to affect subsequent release of glio-modulators. Pharmacological inhibition of the TGFβ pathway partially prevents the synaptogenic effect of murine astrocyte-conditioned media on cortical neurons but abolishes the effect of human astrocyte-conditioned media, suggesting that human astrocytes may rely more heavily upon TGFβ signaling than their rodent counterparts (Diniz et al., 2012). Human astrocytes also display larger cellular diameters with more elaborated and compartmented processes compared with rodent astrocytes (Oberheim et al., 2006). Indeed, while a rodent astrocyte domain can reportedly cover up to 120,000 synapses, a human astrocyte domain can cover up to 2 million synapses, suggesting greater processing complexity in the latter species (Oberheim et al., 2006). These and other species-specific features highlight the potential for human astrocytes to differ from rodent astrocytes in their contributions to brain function as well as brain dysfunction. Human astrocytes thus have important applications in studies of basic brain function, disease modeling and drug discovery.

With the emergence of human pluripotent stem cell (hPSC) technologies, it is now feasible to sustainably generate an array of brain cell types *in vitro*. Notably, numerous studies have shown that glial cells are necessary for the functional maturation of neurons (Nehme et al., 2018; Pfrieger and Barres, 1997; Turovsky et al., 2020). For practical considerations and ease of access, most studies supplement neuronal cultures with rodent astrocytes, and more recently, commercially available primary fetal astrocytes. However, these approaches have significant limitations. As discussed above, rodent astrocytes diverge morphologically, transcriptionally and functionally from human astrocytes and do not allow for the investigation of the effect of human genetic variants and perturbations on biology and disease. Primary fetal astrocytes are not a sustainable resource and generally do not allow for the study of specific human genotypes of interest.

Recognizing the utility of human *in vitro* derived astrocytes, several protocols have been developed including those following a protracted developmental time-course in 3-dimensions (up to 20 mos.) (Sloan et al., 2017) as well as more rapid 2-dimensional protocols (Canals et al., 2018; Hedegaard et al., 2020; Santos et al., 2017; Tcw et al., 2017; Voulgaris et al., 2022). While these protocols produce cells expressing canonical astrocyte markers and are capable of recapitulating key functions such as responding to pro-inflammatory stimuli, a tremendous amount remains to be determined regarding: (i) the precise cell types and cell stages being generated and / or how closely they resemble fetal or adult human astrocytes from primary cultures or post-mortem preparations, (ii) the robustness of protocols across individual hPSC lines, and (iii) their utility for disease modeling applications. Furthermore, most deeply characterized approaches are either lengthy or technically complicated, involving multiple experimental steps, such as purification, replating, and culturing in different formats, rendering such protocols less amenable to implementation across multiple cell lines and manipulations.

Several astrocyte differentiation protocols have recognized the importance of robust NPC generation as an important foundation for efficient astrocyte differentiation (Tcw et al., 2017; Voulgaris et al., 2022). Here, we leveraged previous studies showing that the combination of NGN2 patterning with dual-SMAD and WNT inhibition can direct hPSCs toward diverse neural fates (Limone, 2022; Lin et al., 2021; Nehme et al., 2018; Wells, 2021) as well as one study identifying optimal media conditions to differentiate neural progenitor-like cells (NPCs) into astrocytes (Tcw et al., 2017). Specifically, we used transient NGN2 patterning combined with dual-SMAD inhibition (SB431542, LDN-193189) and WNT inhibition (XAV939) to generate NPCs along a forebrain fate (Nehme et al., 2018; Wells, 2021) followed by maturation in astrocyte media (ScienCell) previously screened for efficient differentiation of hPSC-derived forebrain NPCs into astrocytes (Tcw et al., 2017). This combination resulted in robust generation of astrocytes by 30 days *in vitro*. We then benchmarked our human induced astrocytes (hiAs) against human primary fetal astrocytes (hpAs) using a combination of immunophenotyping, RNA-sequencing and a series of functional assays assessing response to pro-inflammatory stimuli, ATP-induced calcium release and synaptic connectivity upon neuronal co-culture. While single-cell RNA-sequencing (scRNAseq) of *in vitro* derived astrocytes has been limited to 3-dimensional protocols (Rapino, 2022; Sloan et al., 2017), requiring dual transcription factor over-expression approaches (Leng, 2022) or following FACS-purification (Barbar et al., 2020), we used scRNAseq analyses of our rapid 2-dimensional protocol from eight unique parental cell lines to define their molecular signatures. This revealed a striking degree of homogeneity at the single-cell level, and high reproducibility in the differentiated product across multiple parental cell lines. Finally, we generated hiAs in a model of trisomy 21 and recapitulated key features associated with the disease.

Collectively, these analyses establish a rapid, scalable and reproducible differentiation protocol to generate homogeneous human astrocytes with a well-defined molecular signature, with limited interventions, which can be applied to many parental cell lines and specifically for the purposes of disease modeling.

## RESULTS

### Robust and rapid generation of human induced astrocytes (hiAs) from NPCs

To generate hiAs, a doxycycline inducible NGN2 expression construct was introduced into hPSCs through TALEN-mediated stable integration into the AAVS1 safe-harbor locus (Berryer, 2022). Following neomycin selection for iNGN2 integration, iNGN2-hPSCs were briefly switched to N2 medium with doxycycline and small molecule patterning for 48hrs to induce a forebrain NPC-like fate (Nehme et al., 2018) (**Fig. 1A**). After 24hrs of Zeocin selection for NGN2 induction, cells were moved to a commercially available astrocyte medium (ScienCell) for an additional 28+ days. By day 4 post induction, hiAs took on an astrocyte-like morphology, with flat, wide cell bodies beginning to form star-like projections (**Fig. 1A**). By day 30, a vast majority of hiAs expressed canonical astrocyte markers including Aquaporin 4 (AQP4), CD44, Solute Carrier family 1 member 3 (SLC1A3), S100 calcium binding protein B (S100B) and Vimentin (VIM) by immunofluorescence, paralleling results obtained with human primary astrocytes (hpAs)(**Fig. 1B-C**). For example, 99.05% of hiAs and 95.12% of hpAs expressed AQP4, and 93.11% of hiAs and 95.62% of hpAs expressed SLC1A3 (**Fig. 1C**). GFAP showed the most variability between hiAs and hpAs (**Fig. 1C**), consistent with previous studies indicating GFAP is a more variable marker of astrocyte fate *in vitro* (Barbar et al., 2020; Tcw et al., 2017).

**Figure 1:**
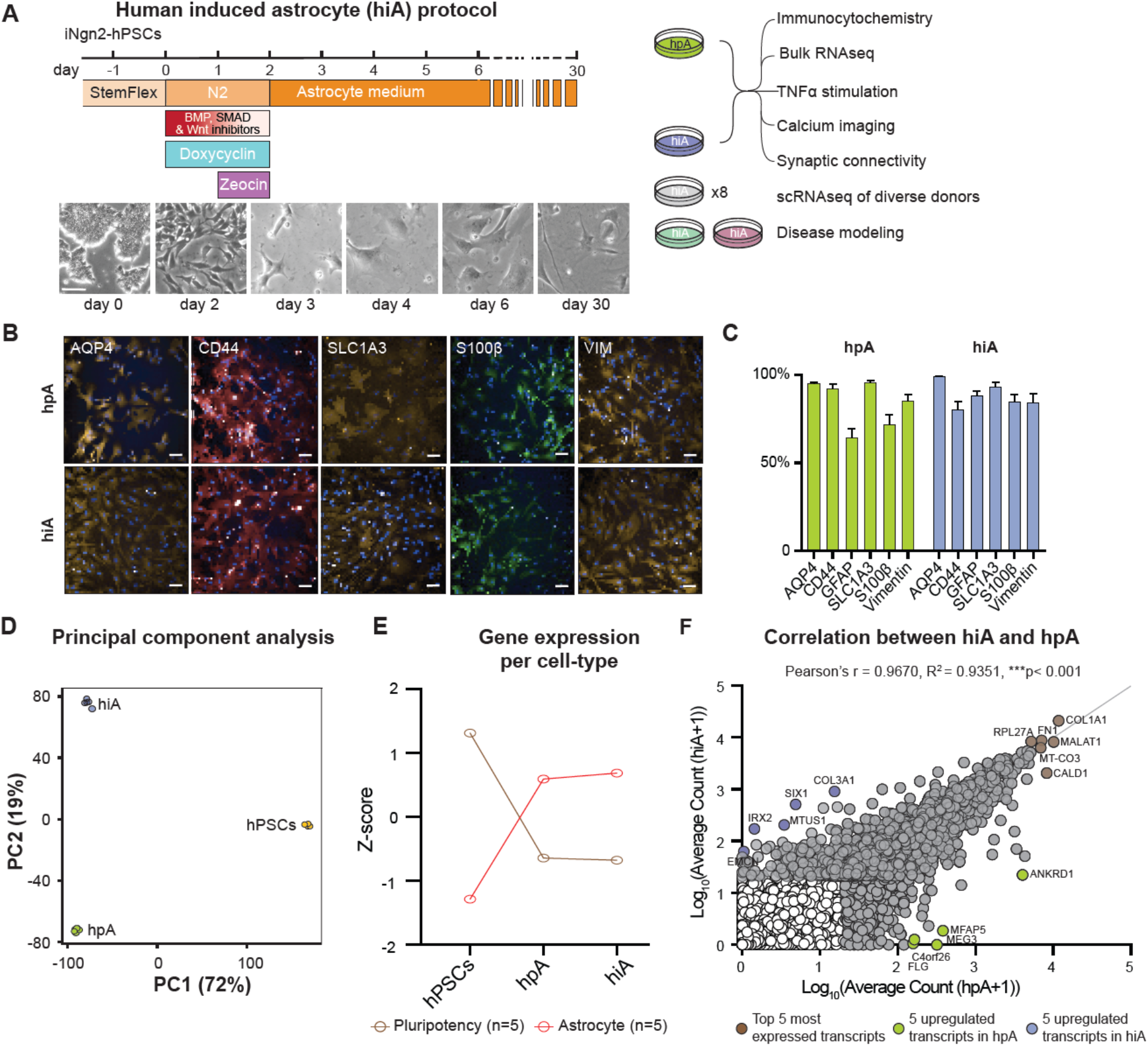
Robust and rapid generation of human induced astrocytes (hiAs) from NPCs. **A**, *Left*, Schematic of the 30-day hiA differentiation protocol with brightfield images over the induction time-course shown below. Scale bar = 100μm. *Right*, Schematic of assays used to assess molecular and functional properties of astrocytes. **B**, Representative immunofluorescence images for AQP4, CD44, SLC1A3, S100b and VIM from human primary astrocytes (top) and human induced astrocytes (bottom). **C**, Quantification of each marker from hiAs and hpAs shown as % of DAPI+ cells. Scale bar = 25μm. **D**, Principal component analysis (PCA) of bulk RNA-seq data comparing hPSCs, hpAs and hiAs. Note that a majority of variance is explained by the comparison between hPSCs and astrocytes. **E**, Gene expression per cell type for hPSCs, hpAs and hiAs using canonical pluripotent and astrocyte related genes. **F**, Scatterplot showing a high positive correlation between hpA and hiA expressed transcripts (TPM >20, in grey). Brown circles highlight the top 5 most expressed transcripts shared between hiAs and hpAs, while green highlights 5 transcripts upregulated in hpAs alone and blue highlights 5 transcripts upregulated in hiAs alone. Pearson’s r = 0.9670, R2 = 0.9351, ***p< 0.001.

To further explore the commitment of hPSCs to an astrocyte identity, we compared the bulk transcriptomic profiles of hiAs with those of hPSCs and hpAs (**Table S1**). Principal component analysis of all three cell types revealed that the vast majority of transcriptomic variance could be explained by the comparison between hPSCs and astrocytes (PC1: 72%) rather than between different astrocyte populations (PC2: 19%) (**Fig. 1D**). Gene expression per cell type revealed a similar trend of decreased pluripotency genes and increased astrocyte-related genes in both hpAs and hiAs (**Fig. 1E**). Finally, we observed a high positive correlation between hpA and hiA expressed transcripts (Pearson’s r = 0.9670; **Fig. 1F**) suggesting a similar landscape in global transcriptome, in contrast with the correlations between hiAs and hPSCs, or hpAs and hPSCs (**Fig. S1**). PANTHER analysis of the top enriched pathways shared by hpAs and hiAs in contrast to hPSCs revealed canonical astrocyte signaling pathways such as VEGF, PDGF, Angiogenesis and Integrin signaling (**Fig. S1**). By contrast, the top enriched pathways unique to hiAs over hpAs or unique to hpAs over hiAs were largely not astrocyte specific (**Fig. S1**). Collectively, these analyses indicate that our hiAs harbor key molecular hallmarks of hpAs.

### hiAs recapitulate key functional features of hpAs

We next sought to establish both cell autonomous and non-cell-autonomous functionality of hiAs as compared with hpAs, including the ability to secrete cytokines in response to pro-inflammatory stimuli, calcium oscillation dynamics in response to ATP and the capacity to promote synaptic development of human *in vitro* derived neurons (hNs). The astrocyte response to pro-inflammatory stimuli is crucial for normal function, with dysfunction of the inflammatory response strongly implicated in disease (Kam et al., 2020; Leal et al., 2013; Liddelow and Barres, 2017; Sofroniew and Vinters, 2010). We therefore challenged our hiAs as well as hpAs with TNFα and quantified subsequent interleukin 6 (IL-6) secretion. Specifically, hpAs and hiAs were treated with either 100ng/mL TNFα or 0.1% BSA control and the harvested supernatant was used for an IL-6 ELISA (**Fig. 2A**). As expected, both hiAs and hpAs showed a robust and significant response to TNFα stimulation compared to BSA control, with hiAs secreting IL-6 at levels slightly above hpAs (p<0.000l; **Fig. 2A**).

**Figure 2:**
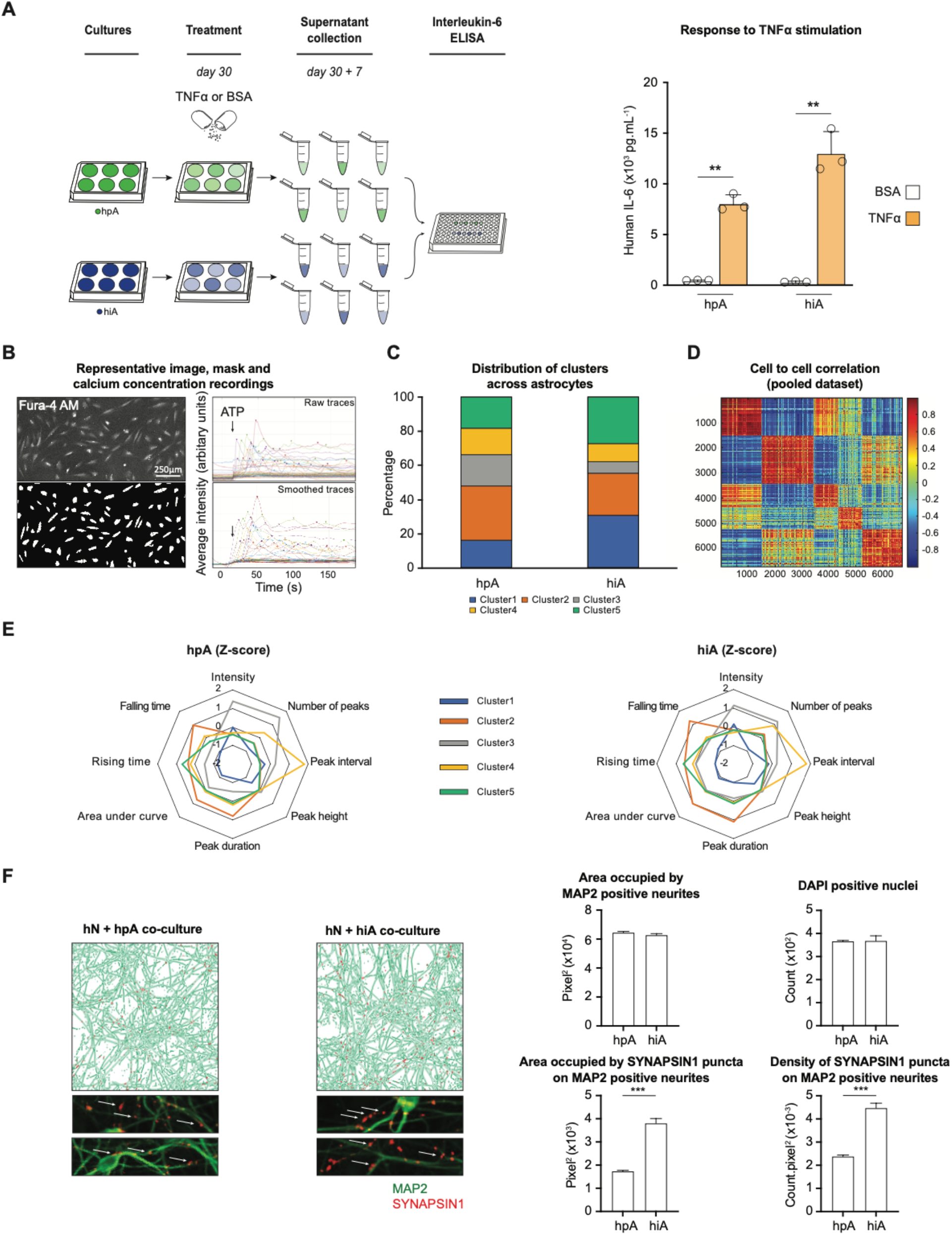
hiAs recapitulate key functional features of hpAs. **A**, *Left*, Schematic of TNFa stimulation experiment. *Right*, Bar graph showing human IL-6 detected from hiAs and hpAs by ELISA in response to 100ng/mL human TNFa (orange) versus 0.1% BSA control (white) treatment. (***p<0.0001; One-way ANOVA with Tukey’s multiple comparisons test: n=3 biological replicates, n=3 technical replicates). **B**, ATP-induced calcium release. Representative Fura-4 image of hiAs (top left), and associated mask (bottom left), raw traces (top right) and smoothed traces (bottom right). Intensity (arbitrary unit) is measured as an average on the surface of each cell. Arrow indicates timing of ATP stimulation. Scale bar = 250μm. **C**, Distribution of each cell population among the identified 5 clusters. Tables of Z-scored features were pulled together, clustered, then split into hpA and hiA. **D**, Correlation heatmap for 6,745 recorded cells. Correlation is computed on the table of Z-scored values for the eight identified features. Cells are sorted by cluster, then within each cluster, are sorted based on their sample of origin, showing that the correlation is strongly dependent on calcium signature but not sample. **E**, Radar plot of the mean Z-scored value taken by the 8 features used for clustering the data. Note that the shape of the typical signature of each cluster is the same for hpAs (left) and hiAs (right). **F**, *Left*, Representative CellProfiler output images of the two conditions (hN + hpA and hN + hiA) and representative pictures of SYNAPSIN1 puncta co-localized with MAP2 positive neurites (arrows point to presynaptic puncta). *Right*, Quantification of the area occupied by MAP2 positive neurites, the area occupied by SYNAPSIN1 puncta colocalized on MAP2 positive neurites, the number of DAPI positive nuclei and the density of SYNAPSIN1 puncta co-localized on MAP2 positive neurites in hN + hpA and hN + hiA co-cultures. (60 wells per condition, ***p<0.0001, unpaired t-test with Welch’s correction).

Astrocytes also display spikes of cytoplasmic calcium concentration as a response to both mechanical, ATP or glutamate stimulation (Allen et al., 2022; Fujii et al., 2017; Turovsky et al., 2020). Here, we compared the response to ATP stimulation of our hiAs and hpAs (**Fig. 2B-E**). Specifically, we used Fura-4 AM dye to image calcium concentration in the cytoplasm, and recorded eight key features including: peak height, number, duration, rising time, falling time, peak interval, area under the curve, and base fluorescence level after stimulation. We recovered the same clusters of typical behavior in response to ATP stimulation in both cell types (**Fig. 2C**). Data from both cell types were then pooled together and k-mean clustered (k=5) (**Fig. 2C-E**). Both hiAs and hpAs were detected in each cluster, with similar distributions overall (**Fig. 2C**). After separating cells based on their origin (hiA versus hpA), we also recovered similar signatures for each cluster (**Fig. 2E**). Notably, individual traces of cells from any given cluster produced the same shape and did not depend on the sample of origin (**Fig. S2**). These data support highly similar calcium dynamics between hiAs and hpAs.

Finally, astrocytes play critical roles in the establishment of synaptic networks, a key non-cell-autonomous function of astrocytes in the human brain. Previous studies have shown that both rodent and human astrocytes can improve the maturation of hNs (Berryer, 2022; Canals et al., 2018; Hedegaard et al., 2020; Nehme et al., 2018). Using an established glutamatergic hN differentiation protocol (Nehme et al., 2018) combined with an automated synaptic quantification platform (Berryer, 2022), we analyzed synaptic development in hN + hiA co-cultures derived from the same parental cell line, as well as hN + hpA co-cultures. These analyses revealed significant increases in the density and the area of presynaptic puncta, identified by SYNAPSIN1 aggregates localized on and along the MAP2-expressing neurites in hiA + hN as compared to hpA + hN co-culture, with no difference in the area occupied by MAP2 positive neurites or the number of DAPI positive nuclei detected. This is consistent with the ability of both hiAs and hpAs to support human synaptic development (**Fig. 2F**).

Collectively, these results demonstrate that, compared to hpAs, our hiAs harbor similar immunocompetence, calcium transients in response to ATP and contributions to the maturation of neuronal networks.

### Single-cell transcriptional profiling of hiAs from multiple donors

Some current hPSC based methods for astrocyte generation require purification steps such as FACS to isolate a specific population of interest given the heterogeneity of cellular differentiation, and variability in the capacity of the differentiation to work effectively across a range of unique cell lines. To better understand the degree of homogeneity and reproducibility in our model, we performed scRNA-seq on 8 unique parental cell lines (**Table S2**). To reduce technical variation, hiAs from each of the 8 donors were differentiated together in a ‘cell village’ as previously described (Mitchell, 2020; Wells, 2021). In brief, hiAs from each of the 8 cell lines were induced and initially differentiated separately, then mixed in equal numbers 25 days post-induction to form a “village” and sequenced 5 days later (at day 30 post-induction). Each individual cell was then assigned to its donor-of-origin using transcribed single nucleotide polymorphisms (SNPs). Uniform Manifold Approximation and Projection (UMAP) showed high homogeneity across individual cell lines, with no cell line clustering distinctly from others using leiden unsupervised clustering (**Fig. 3A**). Our scRNAseq analyses further confirmed that hiAs expressed canonical immature astrocytes markers including *VIM, GJA1, APOE, CD44, SLC1A3*, and *ID3* (**Fig. 3B**). Due to the propensity of *in vitro* differentiation protocols to have high heterogeneity and limited cell-type specificity, we explored whether using our NGN2 over-expression approach would produce oligodendrocyte or neuronal populations among our cells. This was particularly relevant as extended/longer-term NGN2 over-expression can also drive the generation of glutamatergic neurons (Nehme et al., 2018; Zhang et al., 2013). We found that hiAs displayed minimal expression of common genes for both neurons and oligodendrocytes and continued to express limited amounts of stem cell related genes (**Fig. 3C**). To explore the variability across cell lines, we next compared the expression of canonical immature and mature astrocyte markers across donors, as well as global gene expression. Across all 8 cell lines, we observed a high degree of correlation of global expression across donors (Pearson’s r = 0.96-98), highlighting the reproducibility of our method across a range of cell lines (**Fig. 3D**).

**Figure 3:**
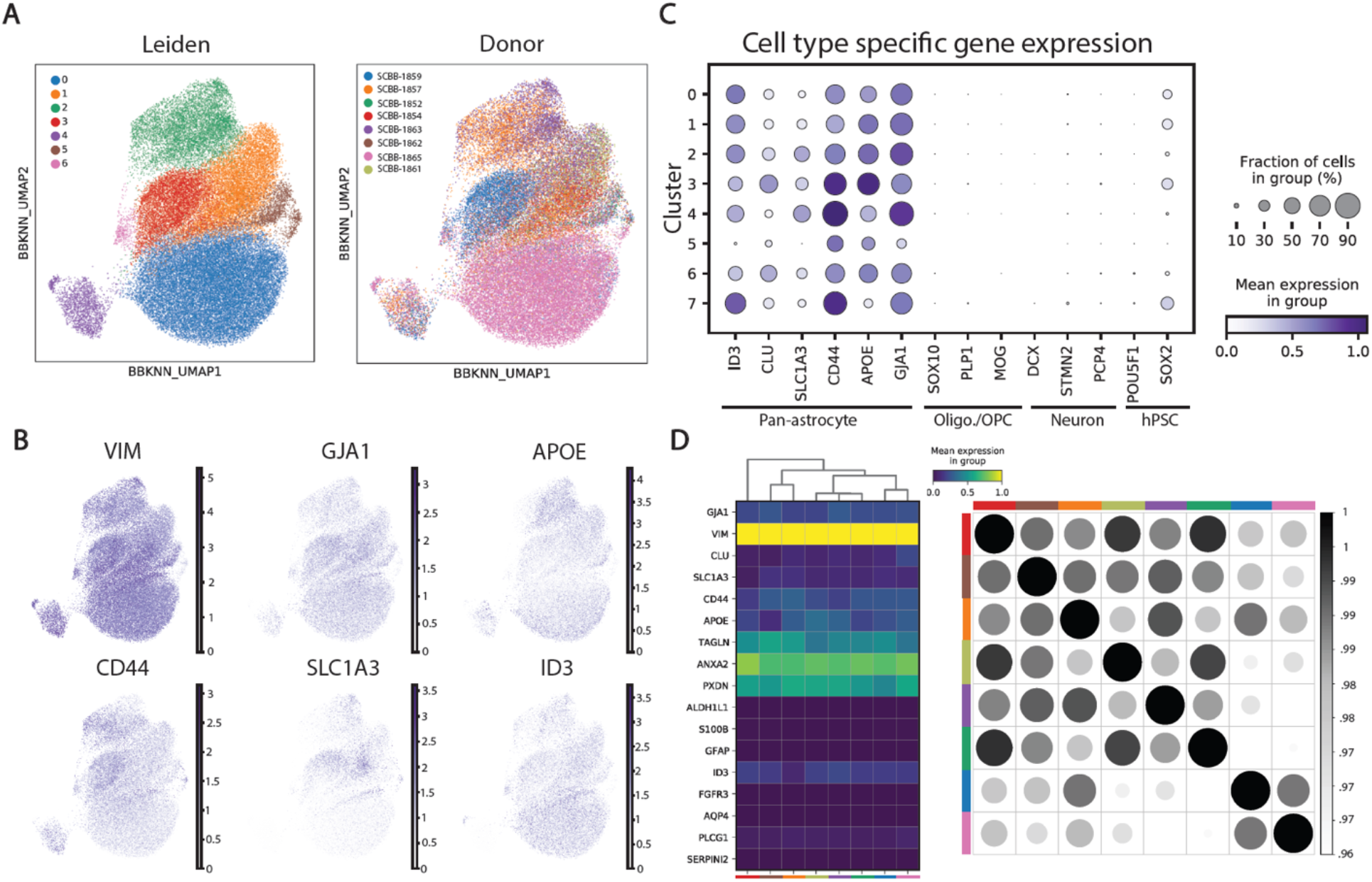
Single-cell transcriptional profiling of hiAs from multiple donors. **A**, *Left*, UMAP projection of all 8 unique parental lines from scRNA experiment labeled by cluster using Leiden unsupervised clustering. *Right*, UMAP projection of all 8 unique parental lines from scRNA experiment labeled by cell line ID. **B**, Feature plot illustrating the distribution of canonical astrocyte markers *VIM, GJA1, APOE, CD44, SLC1A3*, and *ID3* in UMAP space. **C**, Dot plot for markers of astrocytes (*ID3, CLU, SLC1A3, CD44, APOE, GJA1*), oligodendrocytes (*SO×10, PLP1, MOG*), excitatory neurons (*DCX, STMN2, PCP4*), and pluripotent stem cells (*POU5F1, SOX2*). **D**, *Left*, Matrix plot displaying average expression of canonical astrocyte markers by cell line. *Right*, correlation matrix of average expression across all genes between cell lines.

### Comparison of hiAs with *in vitro* and *ex vivo* astrocyte datasets

An important aim of *in vitro* models is to develop cellular substrates which resemble those found in the living human brain. In an effort to understand the similarities and differences among current iPSC-based astrocyte protocols, we compared our hiAs with existing scRNAseq datasets (Barbar et al., 2020; Leng, 2022; Rapino, 2022). We found that UMAP displayed a high degree of similarity across each *in vitro* dataset with the majority of cells clustering together (**Fig. 4A,B**). hiAs showed a high degree of homogeneity, with fewer sub-populations which clustered separately from the majority compared to other datasets (**Fig. 4C**). We next compared the overall correlation of gene expression across iPSC models and found a high level of correlation (*Pearson’s r =* .*65-* .*88*)(**Fig. 4D**). Though global expression across these datasets was similar, we wondered what genes were driving differences between them. We thus performed a differential expression analysis to compare across individual models and found that genes contributing to differences across models were not specific for astrocyte identity or function, with many of the differentially expressed genes including mitochondrial and ribosomal genes **(Fig. 4E, Table S**3). This illustrated that on average, each model resulted in comparable levels of expression of canonical astrocyte markers. While this was true when exploring average expression, there were differences in canonical astrocyte gene expression patterns across clusters, where certain models populated greater proportions (**Table S4**). For example, cluster 9 showed elevated expression of immature astrocyte markers such as *TOP2A, CENPF* and *NUSAP1* relative to other clusters (**Fig. 4F**). As shown in **Fig. 4C**, cluster 9 was predominantly made of cells from Leng et al (Leng, 2022), but only represented ∼ 0.2% of the total population. However, issues such as technical variation and limitations of sequencing depth may impact the detection and direct comparison of specific transcripts across datasets. We therefore examined expression across a large set of genes associated with astrocyte identity and behavior instead of focusing on single genes, creating a metagene score for each cell based on its contribution to expression of a set of genes associated with either astrocyte precursor cells or mature astrocytes as described by Zhang et al, 2016 (Zhang et al., 2016). When we assessed the contribution of each model to these gene sets, we found that all models contributed similar amounts of RNA to both astrocyte precursor cell markers as well as mature astrocyte markers, (**Fig. 4G**). Consistent with the aforementioned findings, hiAs showed lower standard deviations than their counterparts, further illustrating the homogeneity of our cells in their contribution to these specific markers.

**Figure 4:**
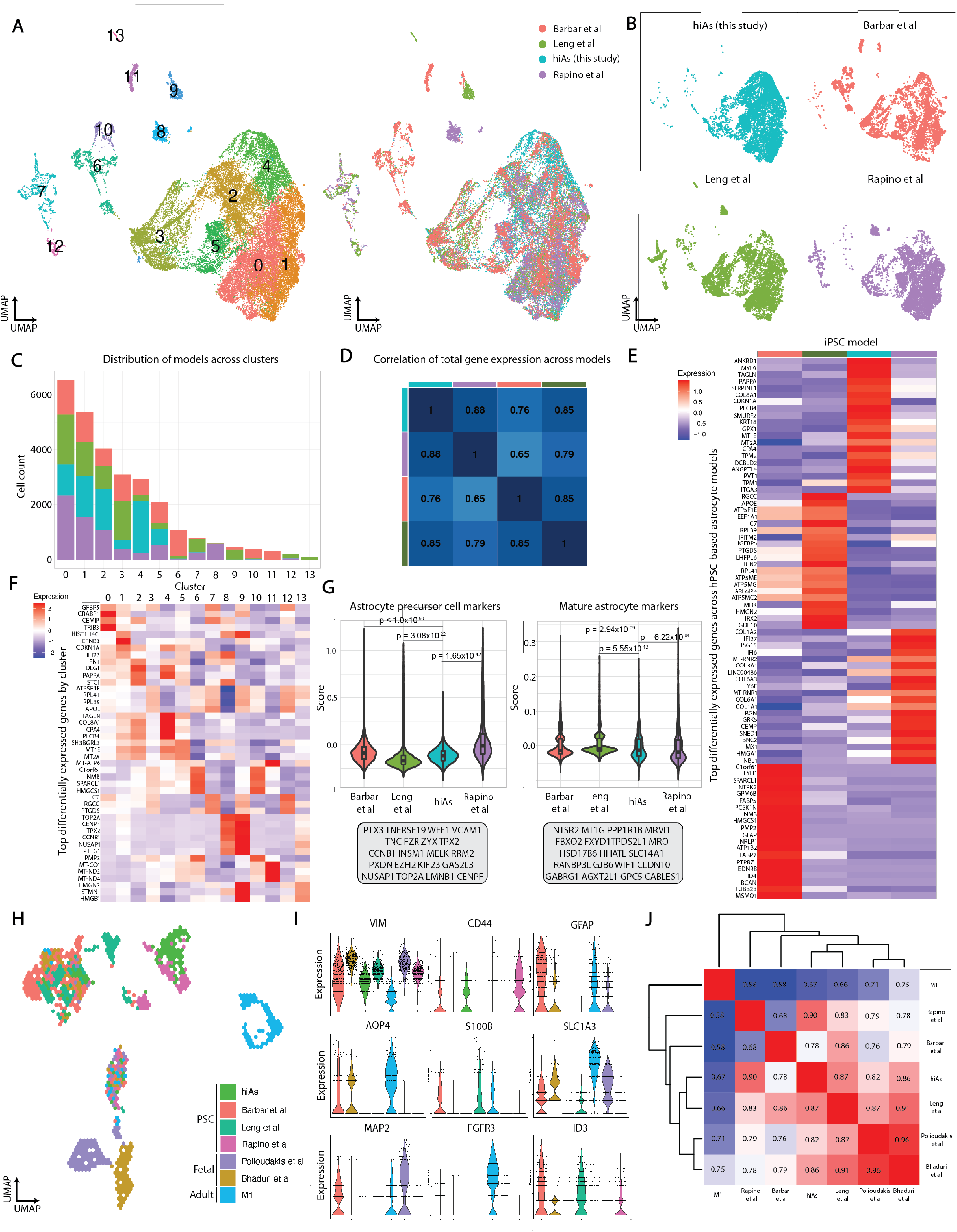
Comparison of hiAs with *in vitro* and *ex vivo* astrocyte datasets. **A**, *Left*, UMAP projection of four iPSC-astrocyte datasets (Barbar et al, Leng et al, Rapino et al and hiAs generated in this study) labeled by cluster using Louvain unsupervised clustering. *Right*, UMAP project of all four iPSC-astrocyte datasets labeled by study. **B**, *Left, top*, UMAP projection only labeling hiAs data. *Left, bottom*, UMAP projection only labeling Leng *et al*. data. *Right, top*, UMAP projection labeling only Barbar *et al*. data. *Right, bottom*, UMAP projection labeling only Rapino *et al*. data. **C**, Distribution of cells from each iPSC-astrocyte dataset by cluster. **D**, Correlation of average gene expression across all iPSC-astrocyte models. **E**, Top 20 differentially expressed genes by iPSC-astrocyte dataset. **F**, Top 4 differentially expressed genes by cluster **G**, *left*, Metagene enrichment score for astrocyte precursor markers across iPSC-astrocyte datasets. p.val = 0.006, ANOVA. *right*, Metagene enrichment score for mature astrocyte markers across iPSC-astrocyte datasets. p.val < 0.005, ANOVA. **H**, UMAP projection of iPSC-astrocyte datasets integrated with fetal and post-mortem human brain data labeled by dataset. **I**, Expression of canonical astrocyte markers (*VIM, CD44, GFAP, AQP4, S100B, SLC1A3, MAP2, FGFR3*, and *ID3*) across iPSC-astrocyte and human brain datasets. **J**, Canonical correlation analysis of iPSC-astrocytes and human brain datasets for average global gene expression.

To understand the *in vivo* fidelity of our system and the other iPSC-based models, we integrated our scRNA-seq data and published datasets with existing data from the fetal human brain (Bhaduri et al., 2021; Polioudakis et al., 2019) and M1 motor cortex from post-mortem adult brain (Network, 2021). UMAP analysis revealed similar clustering between *in vitro* datasets, with some clusters including cells from the two fetal datasets and the M1 atlas used in the analysis (**Fig. 4H**). While some cells generated from each method overlapped with a subset of cells from the human brain tissues, the majority of cells from the fetal and post-mortem datasets still cluster separately (**Fig. 4H**). The expression of some canonical astrocyte genes varied across the brain and stem cell-based datasets (**Fig. 4I**), which may contribute to differences between the human brain datasets and *in vitro* models. For instance, *AQP4, ADH1L1*, and *FGFR3* were more strongly expressed in human adult astrocytes compared to human primary fetal or stem cell derived astrocytes, while *VIM* was robustly expressed across all datasets (**Fig. 4I**). We further compared the average expression across all genes between *in vitro* datasets and the human brain. When looking at the correlation of global gene expression across datasets, hiAs exhibit high correlation with both the fetal brain data, as well as the M1 atlas data (*Pearson’s r =* .*82*, .*86*, .*67, respectively*) (**Fig. 4J**). Additionally, we found that other iPSC-based models exhibited similar correlations to these same human brain datasets, with hiAs showing the strongest correlation to the M1 atlas data relative to other *in vitro* models (*Pearson’s r =* .*67*)(**Fig. 4J**). Based on these analyses, we hypothesized that longer term culture of hiAs might mature them. Interestingly, transcriptomic analysis from hiAs cultured until D60 showed modest changes in gene expression space, and a small increase in correlation to the human brain datasets, suggesting hiAs exhibit a small increase in maturity with additional time in culture, which might be useful for some applications (**Fig. S3**). Overall, our data suggested that hiAs closely resembled other iPSC-derived astrocytes and exhibited strong correlation to relevant human brain datasets, presenting a robust model for investigating astrocyte biology as early as 30 days after differentiation, with a slight improvement from extending time in culture for transcriptional analyses.

### hiAs capture disease phenotypes in model of trisomy 21

We next sought to establish the utility of our hiA protocol for disease modeling, given the high relevance of this application. Specifically, numerous studies have shown deficits in astrocytes due to trisomy 21, which drives Down syndrome (Araujo et al., 2018; Chen et al., 2014; Mizuno et al., 2018; Ponroy Bally et al., 2020; Ponroy Bally and Murai, 2021). For example, using iPSC-derived astrocytes, Chen et al. (2014) found that trisomy 21 drove global transcriptional perturbations and that astrocytes had higher levels of reactive oxygen species and reduced expression of pro-synaptic factors compared to euploid astrocytes (Chen et al., 2014). Also using iPSC-derived astrocytes, Bally et al (2020) identified global transcriptional perturbations due to trisomy 21 coupled with chromatin accessibility analyses, revealing alterations to axon development, extracellular matrix organization and cell adhesion (Ponroy Bally et al., 2020). We therefore generated hiAs from an isogenic pair of commercially available iPSCs with and without trisomy 21 (Weick et al., 2013), referred to here as DS1 (trisomy 21) and DS2U (euploid) hiAs (**Table S5**). As expected, hiAs derived from both euploid and trisomy 21 iPSCs expressed canonical astrocyte markers (**Fig. 5A**), similar to our previous analyses of a control cell line (**Fig. 1B**). Given the high degree of homogeneity observed in our scRNA-seq datasets (**Figs. 3,4**), we then extracted RNA for bulk transcriptional analyses (**Fig. 5C**). As expected, a majority of expressed genes encoded on chromosome 21 (HSA21) were upregulated in trisomy 21 hiAs compared with euploid control hiAs, such as *COL18A1, COL6A, BACE2* and *ADARB1*, with a median fold change of 1.4931 as compared to 0.9623 with the same analysis performed on chromosome 3 (**Fig. 5D; Fig. S4**). Using a p_adj_ threshold of 0.05 and a log_2_FC threshold of +/-2, we also observed global transcriptional perturbations, detecting 1691 significantly upregulated genes and 414 significantly downregulated genes due to trisomy 21 (**Fig. 5E**). GO analyses of the differentially expressed genes revealed biological processes such as cell-cell adhesion, synaptic signaling and neuron projection morphogenesis (**Fig. 5F**), consistent with previous studies of cell-autonomous astrocyte dysfunction as well as deleterious impacts on neuronal and synaptic development in Down syndrome; of note, our analyses captured astrocyte phenotypes also detected in a 160+ day astrocyte differentiation protocol (Ponroy Bally et al., 2020). Examples of individual genes involved in neuron projection morphogenesis and cell-cell adhesion are highlighted in **Fig. 5F, G**. These data support the expected transcriptional dysregulation of hiAs when used in a model of trisomy 21.

**Figure 5:**
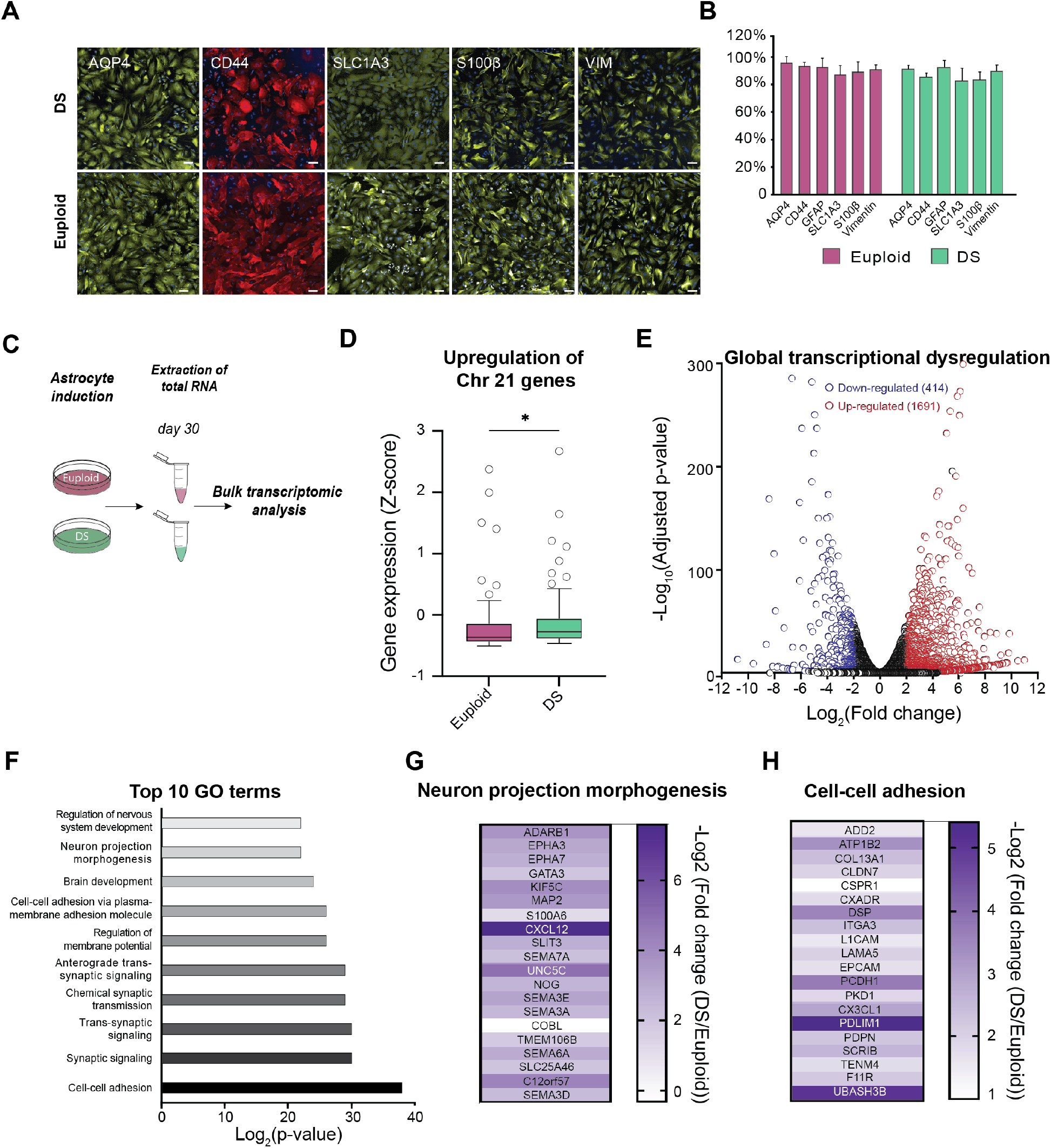
hiAs capture disease phenotypes in model of trisomy 21. **A**, Representative immunofluorescence images for AQP4, CD44, SLC1A3, S100b and VIM from astrocytes derived from DS patient cells (DS1) and astrocytes derived from euploid control cells (DS2U). Scale bar = 100 μm. **B**, Quantification of each marker from DS and euploid astrocytes shown as % of DAPI+ cells. **C**, Schematic of bulk transcriptional analyses for DS and euploid astrocytes. **D**, Gene expression z-scores for 77 genes encoded on chromosome 21 from DS versus euploid control cells. *p=0.0142, Mann Whitney two-tailed t test. **E**, Volcano plot showing wide dysregulation of the transcriptome in DS astrocytes compared to isogenic euploid controls, with a bias towards upregulation. Significantly down-regulated genes are shown in blue and significantly up-regulated genes are shown in red. Log2FC is shown on the x-axis and the −log10 of the adjusted p-value is shown on the y-axis. **F**. The top ten biological processes identified as enriched in differentially expressed genes, as calculated by Gene Ontology (GO) analysis performed via Metascape. The Log2 of the p-value is shown on the x-axis. **G-H**, Heat maps showing examples of genes from the ‘neuron projection morphogenesis’ (left) and ‘cell-cell adhesion’ (right) GO terms identified in (e). Scale shows the −log2FC of DS versus euploid control.

## DISCUSSION

Despite their essentiality for modeling normal brain function as well as dysfunction in disease, glial cell types have lagged somewhat behind neuronal cell types when it comes to hPSC-based differentiation technologies. Protracted astrocyte differentiation protocols as well as those requiring 3-dimensional culture or additional purification steps complicate their utility for many laboratories. Here, we present a simplified, rapid and robust protocol to generate homogeneous astrocytes with detailed molecular and functional benchmarking compatible with many parental cell lines and for disease modeling applications. These astrocytes, generated through a simple protocol driven by transient NGN2 expression, are functional and produce inflammatory responses, elicit calcium signaling, and have promaturational effects on iPSC derived neurons. Further, they are transcriptionally concordant with human primary astrocytes, a common *in vitro* model used to study human astrocyte biology, show a great degree of overlap with existing *in vitro* astrocyte approaches which often require longer culture times or more elaborate interventions to isolate specific cell populations, and display strong correlations with existing human brain datasets. Finally, the hiA cell population is functionally and transcriptionally homogeneous rendering this approach amenable to genetic or pharmacological perturbation screens and circumventing the need for costly single cell sequencing approaches. These features will facilitate the study of human astrocytes for disease modeling as well as drug screening applications.

In the future, we see two key areas to build off from our current hiA differentiation protocol. First, given the high degree of reproducibility across parental cell lines as well as the ability to capture disease-relevant phenotypes and contribute to the maturation of neuronal networks, we see precision co-culture of hiAs with additional brain cell types as a logical next step. Indeed, the advantage of generating each relevant brain cell type separately is the ability to precisely control cell ratios, to generate highly reproducible preparations and to manipulate genotypes or employ genetically diverse parental cell lines to explore cell-type specific effects on network function. We show here that our hiAs can be co-cultured with human neurons and similarly contribute to their development as hpAs, and future studies to incorporate additional brain cell types will facilitate study of a host of biological questions. Second, while hiAs possess key molecular and functional features of hpAs, similar to other iPSC-derived astrocytes, they more closely resemble fetal than adult human astrocytes. It thus is clear more needs to be done to enhance their maturation in order to study the role of human astrocytes in processes beyond development. In this regard, it is interesting to note that additional time in culture only slightly improved hiA maturity, suggesting that additional extrinsic factors may be required for further maturation. Indeed, one possibility is that a more complex co-culture system including neurons or other glial cell types will be required to more accurately recapitulate the *in vivo* environment and further mature the hiAs. It is also possible that additional regulatory networks may need to be activated in order to achieve a more mature astrocyte state.

## MATERIALS AND METHODS

### Stem cell culture

All studies using hPSCs followed institutional IRB and ESCRO guidelines approved by Harvard University. The human ESC line WA01 (H1, XY) and the human iPSC lines DS2U (XY) and DS1 (XY) were commercially obtained from WiCell Research Institute (Thomson et al., 1998; Weick et al., 2013)(www.wicell.org). Additional human iPSC lines are from the Stanley Center Stem Cell Resource (https://sites.google.com/broadinstitute.org/sc-stem-cell-resource/) collection included SCBB-1852 (XY), SCBB-1854 (XX), SCBB-1857 (XY), SCBB-1859 (XY), SCBB-1861 (XY), SCBB-1863 (XX), SCBB-1864 (XY), SCBB-1865 (XY), and available from the NIMH Repository and Genomic Resource (NRGR). Each hPSC line was cultured in feeder-free conditions on Geltrex (15mg/ml, ThermoFisher) in mTeSR1 media with 0.2% Normocin (Invivogen). For routine maintenance, cultured cells underwent daily medium changes and were passaged when reaching 70-80% confluence. Here, new 6-well NUNC plates were coated with Geltrex (15mg/ml) for 1hr at 37C. iPSC colonies were dissociated with Accutase (StemcellTech) for 5-10 min at 37C. After incubation, cells were triturated to remove any excess cells from the plate bottom. Accutase-cell suspensions were added to mTeSR1 medium + 10uM Y-27632 in a 15mL Falcon tube. Cells were centrifuged at 300g x 5 min. The cell pellets were then resuspended in mTeSR1 medium + 10uM Y-27632 and plated across new plates at a desired split ratio (between 1:5 and 1:20). Cells were maintained in a humidified incubator at 37C and 5% CO2. hPSCs between passage 10 and 35 were used in this work.

### Human induced astrocyte (hiA) generation

On day 0, hPSCs were differentiated in N2 medium [500 mL DMEM/F12 (1:1) (Gibco, 11320-033), 5 mL Glutamax (Gibco, 35050-061), 7.5 mL Sucrose (20%, SIGMA, S0389), 5 mL N2 supplement B (StemCell Technologies, 07156)] supplemented with SB431542 (10 μM, Tocris, 1614), XAV939 (2 μM, Stemgent, 04-00046) and LDN-193189 (100 nM, Stemgent, 04-0074) along with doxycycline hyclate (2 μg.mL^−1^, Sigma, D9891) with Y27632 (5 mM, Stemgent 04-0012). Day 1 was a step-down of small molecules, where N2 medium was supplemented with SB431542 (5 μM, Tocris, 1614), XAV939 (1 μM, Stemgent, 04-00046) and LDN-193189 (50 nM, Stemgent, 04-0074) with doxycycline hyclate (2 μg.mL^−1^, Sigma, D9891) and Zeocin (1 μg.mL^−1^, Invitrogen, 46-059). On day 2, N2 medium was supplemented with doxycycline hyclate (2 μg.mL^−1^, Sigma, D9891) and Zeocin (1 μg.mL^−1^, Invitrogen, 46-059). Starting on day 2 human induced neural progenitor-like cells were harvested with Accutase (Innovative Cell Technology, Inc., AT104-500) and re-plated at 15,000 cells.cm^−2^ in Astrocyte Medium (ScienCell, 1801) with Y27632 (5 mM, Stemgent, 04-0012) on geltrex coated plates. Cells were maintained for > 30 days in Astrocyte Medium (ScienCell, 1801).

### Human primary astrocytes (hpAs)

Human primary cortical astrocytes (hpA) were obtained from ScienCell Research Laboratories (1800) and cultured according to the manufacturer’s instructions.

### Immunocytochemistry

Immunofluorescence was performed using an automatic liquid handling dispenser (ApricotDesigns, Personal Pipettor). Cells were washed abundantly in 1x PBS, fixed for 20 minutes in PFA (4%, Electron Microscopy Sciences, 15714-S) plus Sucrose (4%, SIGMA, S0389), washed abundantly in 1x PBS, permeabilized and blocked for 20 minutes in Horse serum (4%, ThermoFisher, 16050114), Triton X-100 (0.3%, SIGMA, T9284) and Glycine (0.1M, SIGMA, G7126) in 1x PBS. Primary antibodies were then applied at 4ºC overnight in 1x PBS supplemented with Horse serum (4%, ThermoFisher, 16050114). The following primary antibodies were used: Rabbit anti-human Aquaporin 4 (1:100, Millipore, AB3594), Rat anti-human CD44 (1:500, ThermoFisher, 14-0441-82), Rabbit anti-human GFAP (1:400, Dako, Z0334), Rabbit anti-human SLC1A3 (1:500, Boster, PA2185), Rabbit anti-human S100b (1:200, Abcam, ab52642) and Rabbit anti-human Vimentin (1:100, Cell Signaling, 3932S).

### TNFɑ stimulation

Human primary and induced astrocytes (> day 30 of differentiation) were seeded at 15,000 cells.cm^−2^ in 96-well plates and incubated 24 hours later with human Tumor Necrosis Factor alpha (100ng.mL^−1^) or 0.1% Bovine Serum Albumin (Sigma Aldrich, A0281-10G) in Astrocyte medium (ScienCell, 1800) for 7 days. The media was replaced with fresh treatment after 4 days of incubation. Media were collected and stored at −80ºC until further processing. IL-6 concentration was measured by ELISA (Abcam, ab229334) in supernatants diluted 1:50 according to the manufacturer’s protocol (Abcam).

### hN generation and synapse quantification

Human induced neuron (hN) generation, astrocyte co-culture and synapse quantification was performed as previously described (Berryer, 2022; Nehme et al., 2018; Zhang et al., 2013). In brief, hNs with stable integration of iNGN2 in the AAVS1 safe-harbor locus were differentiated on day 0 in N2 medium supplemented with SB431542, XAV939 and LDN-193189 along with doxycycline and Y27632. On day 1, N2 medium was supplemented with SB431542, XAV939 and LDN-193189 along with doxycycline and Zeocin. On day 2, N2 medium was supplemented with doxycycline and Zeocin. Starting on day 3, cells were maintained in Neurobasal media (NBM) supplemented with B27, BDNF, CTNF, GDNF and doxycycline. On days 4 and 5, NBM was complemented with floxuridine. On day 6, hNs and hpAs were harvested with Accutase, and plated in NBM using a liquid handling dispenser (Personal Pipetter, ApricotDesigns) in the 60-inner wells of geltrex-coated 96-well plates. Co-cultures were then maintained in NBM until fixation and immunostaining on day 21 of hN differentiation. Immunofluorescence was performed using an automatic liquid handling dispenser (Personal Pipetter, ApricotDesigns) using Rabbit anti-human SYNAPSIN1 (1:1000, Millipore, AB1543) and Chicken anti-human MAP2 (1:1000, Abcam, ab5392) primary antibodies, Goat anti-chicken AlexaFluor 488 (1:1000, ThermoFisher, A21131), Donkey anti-rabbit AlexaFluor 555 (1:1000, ThermoFisher, A31572) secondary antibodies, as well as DAPI (1:5000, ThermoFisher Scientific, D1306) and TrueBlack (1:5000, Biotium, 23007). Images were acquired with a high-content screening confocal microscope (Opera Phenix, PerkinElmer), analyzed with CellProfiler pipelines (https://github.com/mberryer/ALPAQAS).

### Calcium imaging and analysis

Cells were incubated in fura-4AM dye at 2***μ***M for 30 min at 37C. Cells were then washed and imaged in 200 ***μ***L recording solution (125mM NaCl, 2.5mM KCl, 15mM HEPES, 30mM glucose, 1mM MgCl_2_, 3mM CaCl_2_ in water, pH 7.3, mOsm 305). Time lapse videos were acquired at 4X on a Nikon Ti2-E microscope at 2 Hz for 5 mins. Cells were stimulated after 1 min by addition of 200 ***μ***L of ATP at 500***μ*** M in recording solution, generating a final concentration of 250 ***μ***M in the well. Analysis of calcium videos was done using a custom MATLAB script. First, cells were segmented using a watershed algorithm. A table of mean fluorescence per cell across time was generated. Traces were smoothed with a Savitzky-Golay filter with a span of 50 images, aligned on the X-axis by subtracting the minimal value, then the peaks of a minimum height and local prominence of 10, with a minimal distance of 3 seconds between peaks were detected. We then extracted features for each cell: the number of peaks, peak height, interval and duration, rising time and falling time (defined as the time between the peak and the previous/next change of sign in the derivative). Data from all samples (4 wells of primary cells, and 8 wells of induced astrocytes from 2 parental cells, 4 wells each) were Z-scored then pooled together before k-means clustering with k=5. Data were then re-split into 12 tables according to the origin of each cell for statistics. For each cluster, example traces for 60 cells randomly picked from all 12 samples of origin were generated by normalizing all traces between 0 and 255 and generating an 8-bit image (Supplementary fig 2). Scripts are available here https://github.com/lbinan/astrocyteInduction.

### 3’ DGE bulk mRNA-sequencing and analysis of hPSCs, hiAs and hpAs

Four to five biological replicates per cell type of human pluripotent stem cells, human primary cortical astrocytes, and human induced astrocytes (>day 30 of differentiation) were harvested in RLTplus Lysis buffer (Qiagen, 1053393). Total RNA was isolated using the RNeasy micro/mini plus kit (Qiagen, 74034) and stored at −80ºC. Total RNA was quantified using the Qubit 2.0 fluorimetric Assay (Thermo Fisher Scientific). Libraries were prepared from 125 ng of total RNA using a 3’DGE mRNA-seq research grade sequencing service (Next Generation Diagnostic srl) (Xiong et al., 2017) which included library preparation, quality assessment and sequencing on a NovaSeq 6000 sequencing system using a single-end, 100 cycle strategy (Illumina Inc.). The raw data were analyzed by Next Generation Diagnostics srl proprietary 3’DGE mRNA-seq pipeline (v1.0) which involves a cleaning step by quality filtering and trimming, alignment to the reference genome and counting by gene (Anders et al., 2015; Dobin et al., 2013). Differential expression analysis was performed using edgeR (Robinson et al., 2010). Samples were sequenced and analyzed at TIGEM (Pozzuoli, Italy).

### Bulk mRNA-sequencing and analysis of DS1 and DS2U hiAs

Two biological replicates of DS1 and two biological replicates DS2U hiAs were harvested in RLTplus Lysis buffer (Qiagen, 1053393). Total RNA was isolated using the RNeasy micro/mini plus kit (Qiagen, 74034). Libraries were prepared using Roche Kapa mRNA HyperPrep strand specific sample preparation kits from 200ng of purified total RNA according to the manufacturer’s protocol using a Beckman Coulter Biomek i7. The finished dsDNA libraries were quantified by Qubit fluorometer and Agilent TapeStation 4200. Uniquely dual indexed libraries were pooled in equimolar ratio and subjected to shallow sequencing on an Illumina MiSeq to evaluate library quality and pooling balance. The final pool was sequenced on an Illumina NovaSeq 6000 targeting 30 million 100bp read pairs per library. Sequenced reads were aligned to the UCSC hg19 reference genome assembly and gene counts were quantified using STAR (v2.7.3a) (Dobin et al., 2013). Differential gene expression testing was performed by DESeq2 (v1.22.1) (Love et al., 2014). RNAseq analysis was performed using the VIPER snakemake pipeline (Cornwell et al., 2018). Library preparation, Illumina sequencing and VIPER workflow were performed by the Dana-Farber Cancer Institute Molecular Biology Core Facilities. Gene Ontology (GO) analysis of differentially expressed genes was performed using Metascape (Zhou et al., 2019).

### scRNA-sequencing and donor assignment

For single-cell analyses, cells were harvested and prepared with 10X Chromium Single Cell 3’ Reagents V3 and sequenced on a NovaSeq 6000 (Illumina) using a S2 flow cell at 2 × 100bp. Raw sequence files were then aligned and prepared following the Drop-seq workflow (Macosko et al., 2015). Human reads were aligned to GRCh18 and filtered for high quality mapped reads (MQ 10). In order to identify donor identity of each droplet, variants were filtered through several quality controls as described previously to be included in the VCF files (Wells, 2021), with the goal of only using sites that unambiguously and unequivocally can be detected as A/T or G/C. Once both the sequenced single-cell libraries and VCF reference files are filtered and QC’ed, the Dropulation algorithm is run. Dropulation analyzes each droplet, or cell, independently and for each cell generates a number representing the likely provenance of each droplet from one donor. Each variant site is assigned a probability score for a given allele in the sequenced unique molecular identifier (UMI) calculated as the probability of the base observed compared to expected based, and 1 – probability that those reads disagree with the base sequenced. Donor identity is then assigned as the computed diploid likelihood at each UMI summed up across all sites (Wells, 2021).

### scRNAseq analysis of villages and integrated datasets

Gene by cell matrices from hiA villages were built from separate runs of 10X Chromium Single Cell 3’ Reagents V3 as described above. Cells with less than 200 genes and more than 15% mitochondrial RNA were trimmed away from downstream analyses. SNN graphs were computed using batch-balanced k-nearest neighbors (BBKNN) to remove batch effects across 10X reactions (Polanski et al., 2020). Leiden clustering was performed across resolutions in BBKNN space (0.2,0.4,0.6) and then visualized to determine the dimensionality of the data. LEIDEN_BBKNN_0.2 was used for the downstream analysis. For the metagene analysis, summed expression of gene sets for astrocyte precursor markers and mature astrocyte markers were divided by a random control set of 500 genes. For the analyses where we integrated existing iPSC-astrocyte data as well as human brain data, raw matrices were loaded in Seurat v4.0.1. The same parameters were applied as above, excluding cells with fewer than 200 genes and greater than 15% mitochondrial gene expression. Given the technical variability across datasets, and cell sources, we first computed a new count matrix using SCTransform (Hafemeister and Satija, 2019). Next, the transformed data was integrated using linked inference of genomic experimental relationships (LIGER) (Welch et al., 2019). Downstream analytical steps were performed using Seurat v4.0.1 basic functions.

### Data, resource and code availability

Bulk RNA-seq data are provided in Tables S1 and S5. Single cell RNA-seq data is available via the Broad Institute Single Cell Portal (https://singlecell.broadinstitute.org/single_cell/study/SCP1972/berryer-tegtmeyer-et-al-hpsc-derived-astrocytes#study-summary). All other data is available in the manuscript or the supplemental materials. Engineered cell lines are available upon request to the corresponding authors and following appropriate institutional guidelines for their use and distribution.

## ACKNOWLEDGEMENTS

We thank members of the Barrett and Nehme labs for insightful discussions and critical reading of the manuscript. We appreciate the analytical support from Zach Herbert at the Dana-Farber Cancer Institute Molecular Biology Core Facilities and from TIGEM (Pozzuoli, Italy). This work was supported by a Broad*next*10 grant and R01HD101534 to L.E.B., U01MH115727 to S.A.M. and R.N., R21MH120423 and RF1MH121289 to S.L.F, as well as support from the Stanley Center for Psychiatric Research.

## AUTHOR CONTRIBUTIONS

M.H.B., M.T., R.N. and L.E.B. conceived the project and wrote and edited the manuscript, M.T. performed astrocyte differentiations for the brightfield imaging time-course and scRNAseq experiments and performed analyses with support from D.M. and S.A.M., M.H.B. generated and optimized the hiA protocol, performed immunocytochemistry, bulk RNA-seq, ELISA, and synaptic connectivity experiments and analyzed relevant data, V.V. and L.B. performed the astrocyte calcium release experiments, and L.B. performed the analysis with supervision from S.L.F., O.P. and A.N. assisted with bulk RNA-seq analysis, J.K., E.C. and D.T. assisted with astrocyte production and immunocytochemistry. Astrocyte data from Rapino et al was obtained courtesy of F.R. and L.L.R. All authors discussed the results and edited the manuscript. S.A.M., R.N. and L.E.B. secured funding and oversaw experiments.

## SUPPLEMENTAL INFORMATION

**Fig. S1:**
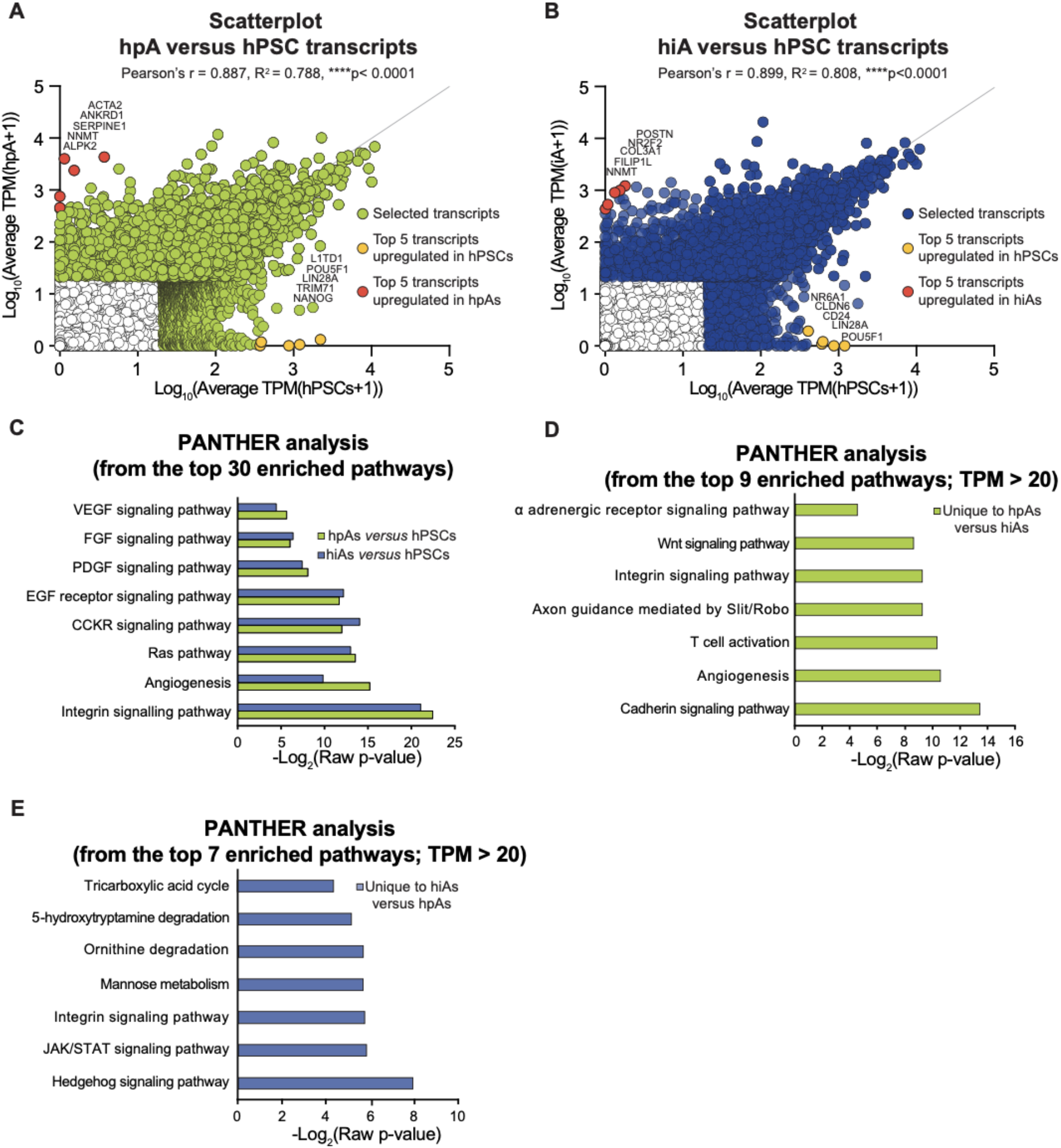
Transcriptomic comparison between hpAs, hiAs and hPSCs. **A**, Scatterplot of the correlation between hPSCs and hpA expressed transcripts. Pearson’s r = 0.887, R2 = 0.788, ****p< 0.0001. **B**, Scatterplot of the correlation between hPSCs and hiA expressed transcripts. Pearson’s r = 0.899, R2 = 0.808, ****p<0.0001. **C**, Biological pathways enriched in differentially expressed genes from hpAs (green) and hiAs (blue) in comparison to hPSCs using PANTHER analysis. **D**, Biological pathways enriched in hpAs in comparison to hiAs using PANTHER analysis. **E**, Biological pathways enriched in hiAs in comparison to hpAs using PANTHER analysis.

**Fig. S2:**
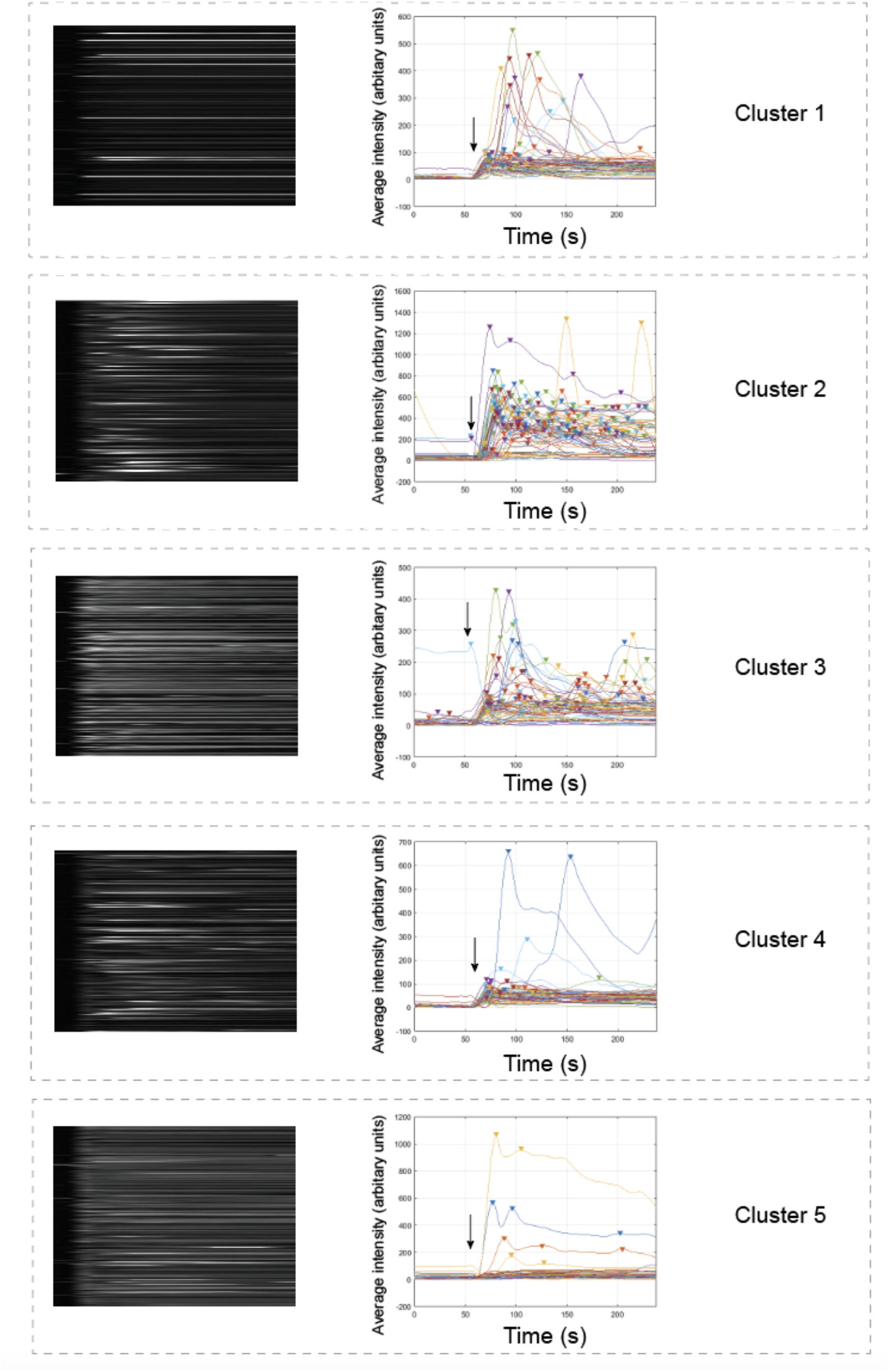
Examples of raw traces and calcium oscillations following ATP addition for typical behavior (cluster). Arrows indicate ATP addition. Plots show the traces of 60 randomly chosen cells from all 12 samples of origin.

**Fig. S3:**
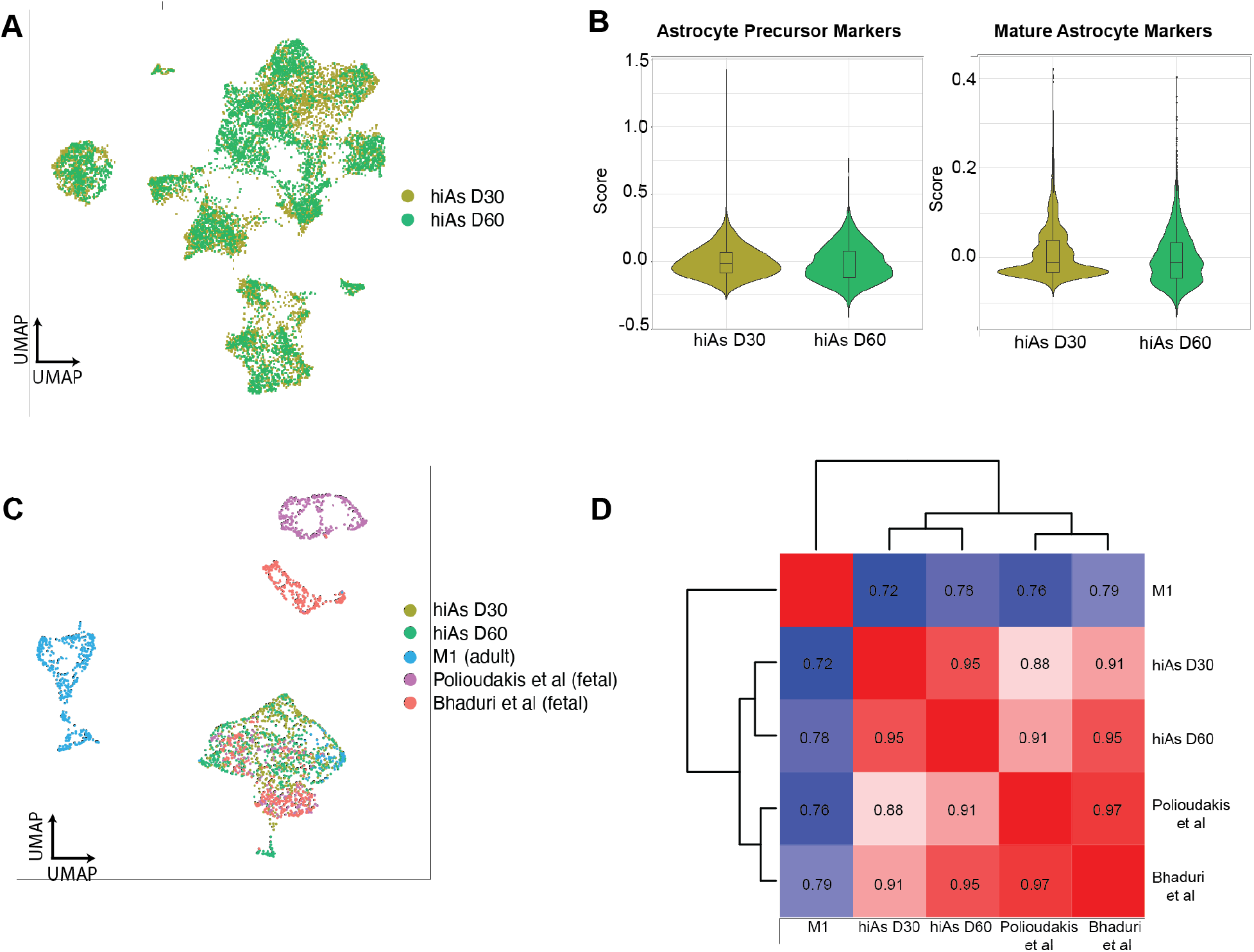
Comparison of D30 and D60 hiAs. **A**, UMAP projection of D30 and D60 hiAs labeled by dataset. **B**, Metagene enrichment score for astrocyte precursor markers and mature astrocyte markers between D30 and D60 hiAs. **C**, UMAP projection of D30 and D60 hiAs integrated with human brain reference datasets. **D**, Canonical correlation analysis between D30 and D60 hiAs integrated with human brain reference datasets.

**Fig. S4:**
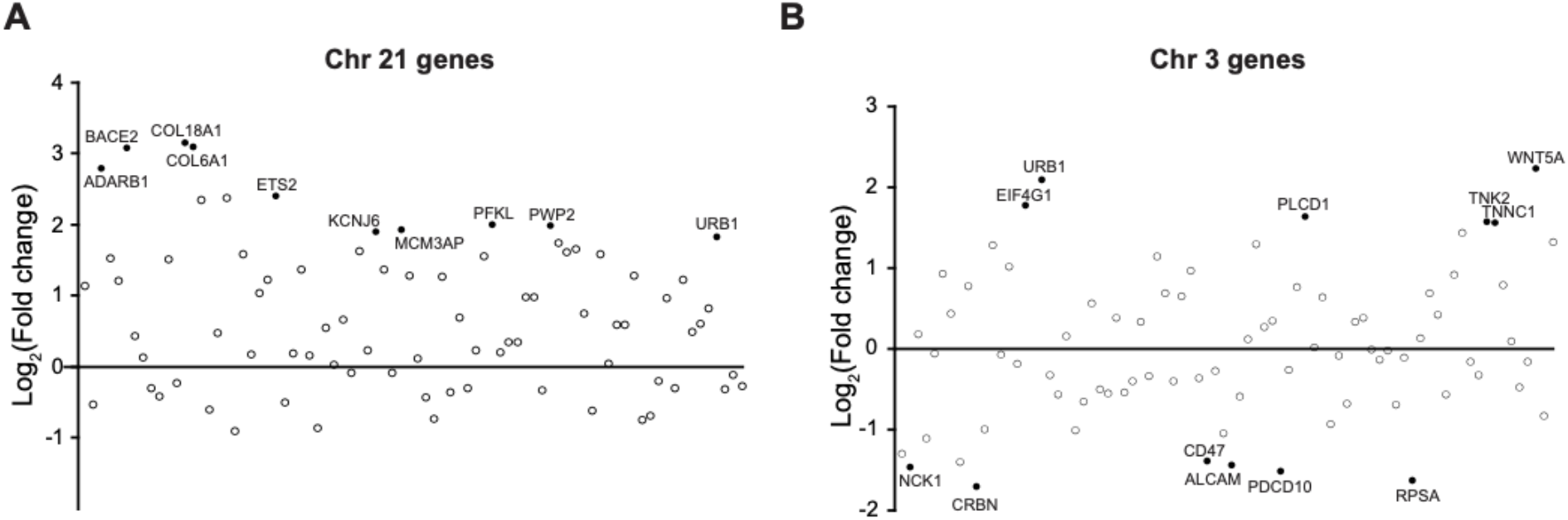
Differentially expressed genes per chromosome from DS2U versus DS1 hiAs. Plots showing Log2FC for genes encoded on chromosome 21 (**A**) and chromosome 3 (**B**) comparing DS1 versus DS2U hiAs. Note the expected upregulation of genes on chromosome 21 as compared with chromosome 3.

**Table S1: Bulk transcriptional analyses of hpAs and hiAs**. Related to Figure 1 and S1.

**Table S2: Single-cell transcriptional analyses**. Per donor average expression from hiA village data. Related to Figure 3.

**Table S4: Single-cell transcriptional analyses**. Differential expression by study in iPSC-astrocyte integrated data. Related to Figure 4.

**Table S3: Single-cell transcriptional analyses**. Differential expression by cluster in iPSC-astrocyte integrated data. Related to Figure 4.

**Table S5: Bulk transcriptional analyses of DS1 and DS2U hiAs**. Related to Figure 5 and S4.

